# Transient disruption of the inferior parietal lobule impairs action mindreading

**DOI:** 10.1101/862284

**Authors:** Jean-François Patri, Atesh Koul, Marco Soriano, Martina Valente, Alessio Avenanti, Andrea Cavallo, Stefano Panzeri, Cristina Becchio

## Abstract

Although it is well established that fronto-parietal regions are active during action observation, whether they play a causal role in the ability to “mindread” others’ actions remains controversial. In experiments reported here, we combined offline continuous theta-burst stimulation (cTBS) with computational modeling to reveal single-trial computations in the inferior parietal lobule (IPL) and inferior frontal gyrus (IFG). Participants received cTBS over the left IPL and IFG, in separate sessions, before completing an intention discrimination task or a kinematic discrimination task unrelated to intention. We found that transient disruption of activity of the IPL, but not the IFG, specifically impaired the observer’s ability to judge intention from movement kinematics. Kinematic discrimination unrelated to intention, in contrast, was largely unaffected. Computational analyses revealed that IPL cTBS did not impair the ability to ‘see’ changes in movement kinematics, nor did it alter the weight given to informative versus non-informative kinematic features. Rather, it selectively impaired the ability to link variations in informative features to the correct intention. These results provide the first causal evidence that IPL maps kinematics to intentions.

## Introduction

When watching others in action, we readily infer their intentions from subtle changes in the way they move. Theoretical work^1–5^ and related experimental findings (e.g.,^6–9^) suggest that this ability to ‘mindread’ the action of others is mediated by the fronto-parietal action observation network. Despite two decades of research, however, the specific neural computations involved in the ability to read the intention of an observed action remain unclear and causally untested.

A major difficulty in studying action mindreading is the ever-changing nature of movement kinematics^10,11^. Movement is “repetition without repetition”^12^. Averaging across repeats of nominally identical, but actually different motor acts, as done in standard trial-averaged analyses, can obscure how intention information is encoded in trial-to-trial variations in movement kinematics^13^. More importantly, the brain does not operate according to an average response over averaged kinematics. Real-world action mindreading requires real-time readout of intention-information encoded within a specific motor act. Thus, studying action mindreading with single-trial resolution is critical for understanding how intention readout maps to the multiplicity and variability of kinematic patterns.

Here, we developed a novel analysis framework to capture intention mapping at the single-trial level. This framework was inspired by recent advances in our understanding of how sensory information encoded in a neural population is read out to inform single-trial behavioral choice^14,15^. In this study, we extended this approach to investigate neural computations performed in the left inferior parietal lobule (IPL), a core region of the action observation network, and explore the hypothesis that neural computations of the left IPL play a causal role in action mindreading.

Activity in IPL is related to the coding of intention in both monkeys and humans^3,6,7,16–18^. In monkeys, the PFG area, found on the convex of the IPL, contains visuo-motor neurons whose activity is modulated by the inferred intention of an observed act (e.g., grasp-to-eat)^6,7^. In humans, intentions inferred from a set of observed actions can be decoded using spatial patterns of activity in the left IPL^16^. These results suggest that the left IPL contains information about intentions of observed actions. However, the level of causal inference afforded by measurement experiments is limited^19,20^. This is because, in the absence of perturbation of neural activity, the relationship between neural information and behavior remains correlational. Moreover, simple observation of neural activity cannot determine what features of neural representations are read out by other regions and what neural computations affect downstream processing^21–23^. The contribution of IPL to action mindreading, both in terms of function – *does IPL plays a causal role in action mindreading?* – and content – *what and how does IPL compute?* – remains therefore largely undefined^24^.

To investigate these questions, we applied continuous theta-burst transcranial magnetic stimulation (cTBS) to reversibly reduce cortical excitability in the left IPL and the left inferior frontal gyrus (IFG), another key region of the action observation network^25–29^. We investigated how transient disruption of activity in these regions influence the observer’s readout computations involved in extracting intention-related information from movement kinematics.

Single-trial analyses combined with a set of task manipulations revealed that disruption of activity in the left IPL, but not the left IFG, selectively impair an observer’s ability to interpret the intentional significance of discriminative kinematic features.

## Results

### Causal contribution of IPL to intention discrimination

To perturb the fronto-parietal action observation network, we used cTBS^30–32^. This protocol delivered offline for 40 s decreases cortical excitability up to 50 min^33,34^. In three separate sessions, participants either received no cTBS or Magnetic Resonance Imaging (MRI)-guided cTBS to the left IPL or left IFG before completing a two-alternative, forced-choice (2AFC) discrimination of intention (Fig. 1a-c).

**Fig. 1.**
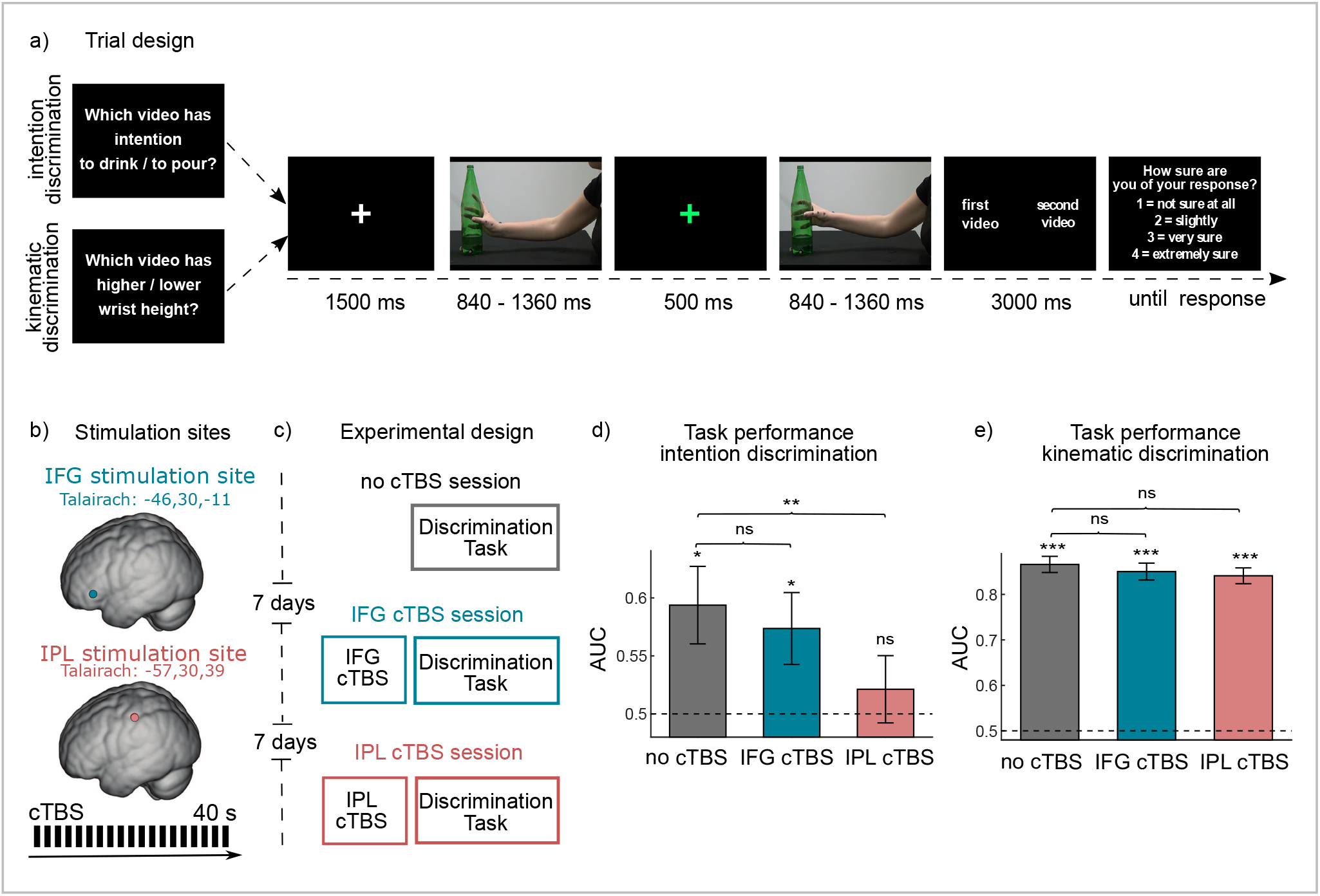
Experimental design and behavioral results. **a** Trial structure of the discrimination tasks. **b** Stimulation sites and MRI-guided cTBS protocol. **c** Sketch of the experimental design. **d-e** Task performance quantified as AUC in the intention discrimination task and in the kinematic discrimination task. Error bars indicate standard error of the mean.

To capture variability in movement kinematics, we employed a dataset of 512 reach-to-grasp acts obtained by simultaneously filming and motion-tracking the kinematics of 17 naïve participants reaching toward and grasping a bottle with the intent to drink or to pour. We extracted 16 kinematic variables over four time periods (0-25%, 25-50%, 50-75%, and 75-100% of movement duration) and selected 60 representative grasping acts for each intention to be used as video stimuli for the intention discrimination task. Each trial displayed two reach-to-grasp acts in two consecutive temporal intervals: one interval contained a grasp-to-pour act, the other interval a grasp-to-drink act. Participants were required to indicate, on each trial, the interval displaying the reach-to-grasp performed with the intent to drink (or to pour; see Methods), and, at the end of the trial, to rate the confidence of their choice (Fig. 1a). We used participants’ responses and confidence ratings to determine points on an empirical receiver operating characteristic (ROC) curve. Average performance over trials, as determined by the area under the ROC curve (AUC), revealed a specific influence of IPL cTBS on the ability to discriminate intention. Specifically, whereas IFG cTBS had no significant effect on task performance, cTBS to IPL significantly impaired ability to discriminate intention (Fig. 1d).

### Functional selectivity of IPL causal contribution

To determine the selectivity of IPL contribution to action mindreading, we repeated our measurements in a new cohort of participants carrying out a 2AFC kinematic discrimination task unrelated to intention. Stimuli and task parameters were identical to that of the intention discrimination task except that participants were required to discriminate differences in the peak wrist height of the observed acts. As shown in Fig. 1e, cTBS to the left IPL (or to the left IFG) did not significantly influence performance in this kinematic discrimination task. Restricting the analysis to less discriminable trials yielded a similar pattern of results, suggesting that the lack of cTBS effects in the kinematic discrimination was not related to the relative ease of the task (see Supplementary Data 1). Collectively, these analyses suggest that observers retain the ability to process changes in movement kinematics following IPL cTBS. Thus, the decrease in task performance in the intention discrimination task post IPL cTBS is not attributable to an inability to ‘see’ changes in movement kinematics *per se*. Taken together, these results provide causal evidence for the selectivity of the effect of IPL cTBS to action mindreading.

### Using logistic regression to relate intention encoding and readout at the single-trial level

There are at least two ways in which transient disruption of activity in left IPL could selectively impair action mindreading. First, following IPL cTBS, observers may be capable of processing changes in movement kinematics (as demonstrated by performance in the kinematic discrimination task), but not be able to use such changes to judge intention. Statistically, this would be reflected in a decrease of the sensitivity of intention readout to single-trial variations in movement kinematics. Alternatively, observers may use changes in movement kinematics to judge intention but be unable to link such changes to the correct intention; that is, access to the intentional significance of the observed differences is hindered. These hypotheses make distinct predictions regarding how the transient disruption of IPL influences trial-wise dependencies between intention encoding and readout. To distinguish between these alternatives, we employed logistic regression to analyze how intention-related information encoded in movement kinematics is read out at the single-trial level^22,35^.

### Encoding of intention-related information

To obtain a measure of intention-related information encoded in grasping kinematics with single-trial resolution, we developed an encoding model based on logistic regression^36^. This model incorporated single-trial changes in movement kinematics as predictors of the intention of the observed reach-to-grasp acts. For each trial, variations in movement kinematics were quantified as a 64-dimensional vector (shortened hereafter to “single-trial kinematic vector”) of the differences between the first and the second reach-to-grasp act for each kinematic feature (16 kinematic variables x 4 time periods). The encoding model computed the probability of the first interval in each trial to display a grasp-to-drink act (and thus of the second interval to display a grasp-to-pour act) as a sigmoid transformation of the linear combination of the features of the kinematic vector for that trial (Fig. 2c, d). Fig. 2e shows a sketch of the encoding model in a hypothetical, simplified kinematic space spanning only two kinematic features. The encoding boundary defines the border that best separates the kinematic patterns of the two intentions. The encoding vector, orthogonal to the encoding boundary, indicates the information axis along which changes in kinematics maximally discriminate between intentions. Single-trial kinematic vectors were classified as ‘to pour’ or ‘to drink’ depending on which side of the boundary they fell or, equivalently, according to the angle they formed with the encoding vector. Since in our convention the encoding vector pointed towards ‘to drink,’ 0-90° encoding angles indicated ‘to drink,’ whereas 90-180° encoding angles indicated ‘to pour.’

**Fig. 2.**
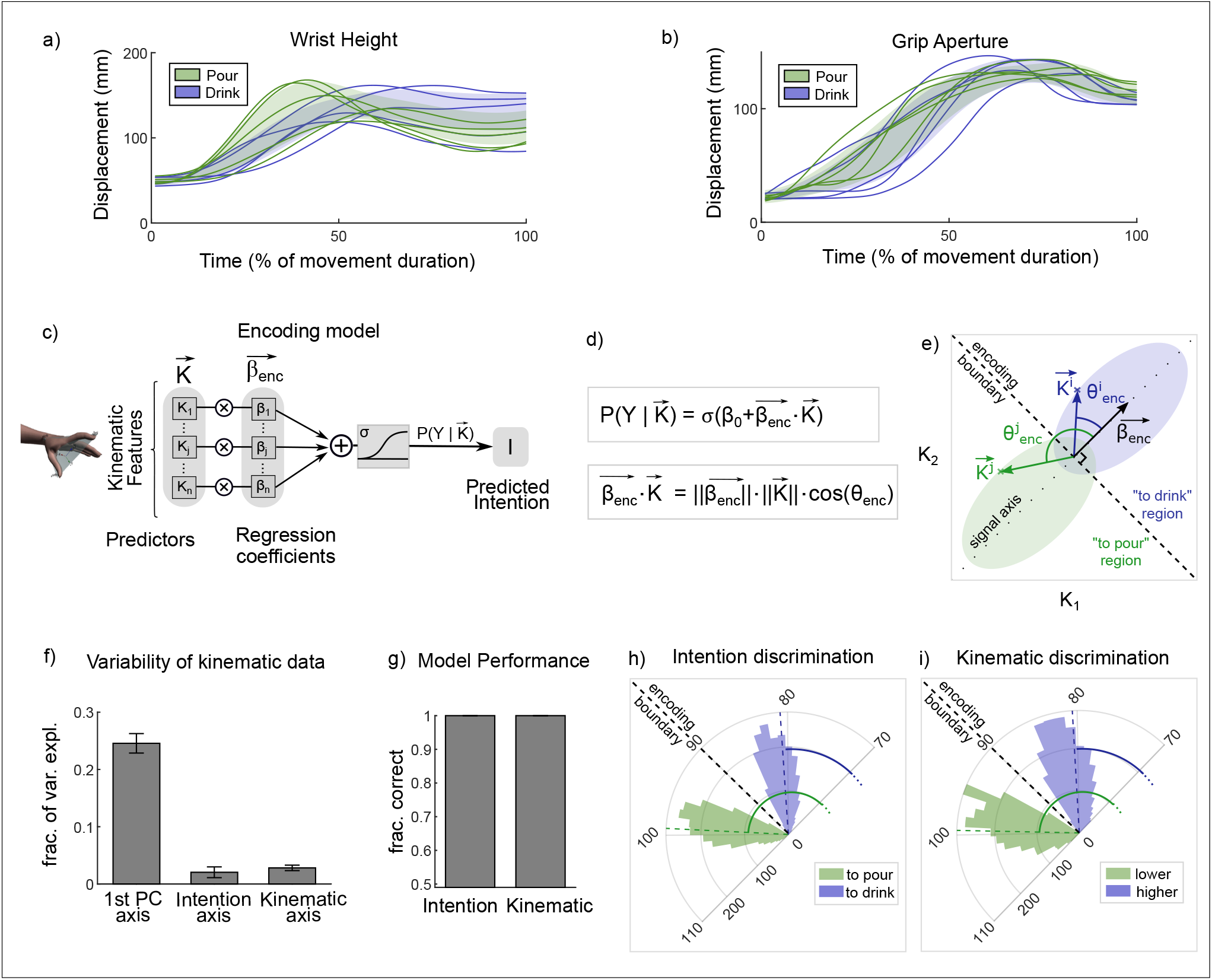
Encoding of task discriminant information in movement kinematics. **a-b** Time course of wrist height and grip aperture for reach-to-grasp acts performed with the intent to drink and to pour. Colored curves display representative trajectories for each intention, colored areas display one standard deviation across executed trials. **c-d** Schematic of the encoding model. Block diagram representation (c) and equation (d) of the logistic regression used to quantify intention information in movement kinematics. **e** Sketch of the encoding model in a simplified kinematic space spanning only two kinematic features. The two elliptic regions represent the intention conditional probability distributions of the two features. The encoding boundary optimally separates the kinematics patterns into ‘to drink’ and ‘to pour’ regions. The encoding vector 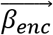 indicates the maximally discriminative axis. The angle *θ_enc_* between the encoding vector and the single-trial kinematic vector 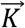 can be used to classify single trial according to intention. Two different single-trial kinematic vectors with index *i* and *j* are shown. **f** Fraction of variance explained across executed acts computed along three different kinematic axes (the PC1 axis of largest variance, the axis of largest intention discrimination, and the axis of largest wrist height discrimination). **g** Performance of encoding models, quantified as the fraction of correctly predicted trials. **h-i** Distribution of encoding angles across trials in the intention discrimination task (h) and in the kinematic discrimination task (i). For graphical representation, the 70-110° angle range is expanded to a semi-circle. Error bars indicate standard error of the mean.

Kinematic features were variable across individual trials, with only a small amount of variance (about 3%) aligned along the encoding vector and thus available for intention discrimination (Fig. 2f). Despite the small amount of intention-related signal hidden within the highly variable kinematic data, our encoding model was able to decode intention with 100% accuracy (Fig. 2g, h). This indicates that intention-related variation in grasping kinematics, although small, is nevertheless sufficient to specify intention-information in each trial. As expected by task design, a 100% decoding accuracy was also observed for the kinematic discrimination task (Fig. 2g, i).

### Readout of intention-related information

Having quantified intention-specifying information encoded in single-trial kinematics, we next asked how human observers read out such information. To assess this, we developed a readout model that predicted the probability of intention choice on each trial as a sigmoid transformation of the linear combination of the features of the kinematic vector for that trial (Fig. 3a-c). Regression coefficients were estimated separately for each subject for no cTBS, IPL cTBS, and IFG cTBS sessions. Given that IFG cTBS did not affect behavioral performance, single-trial analyses for IFG cTBS are reported as Supplementary Data 6.

**Fig. 3.**
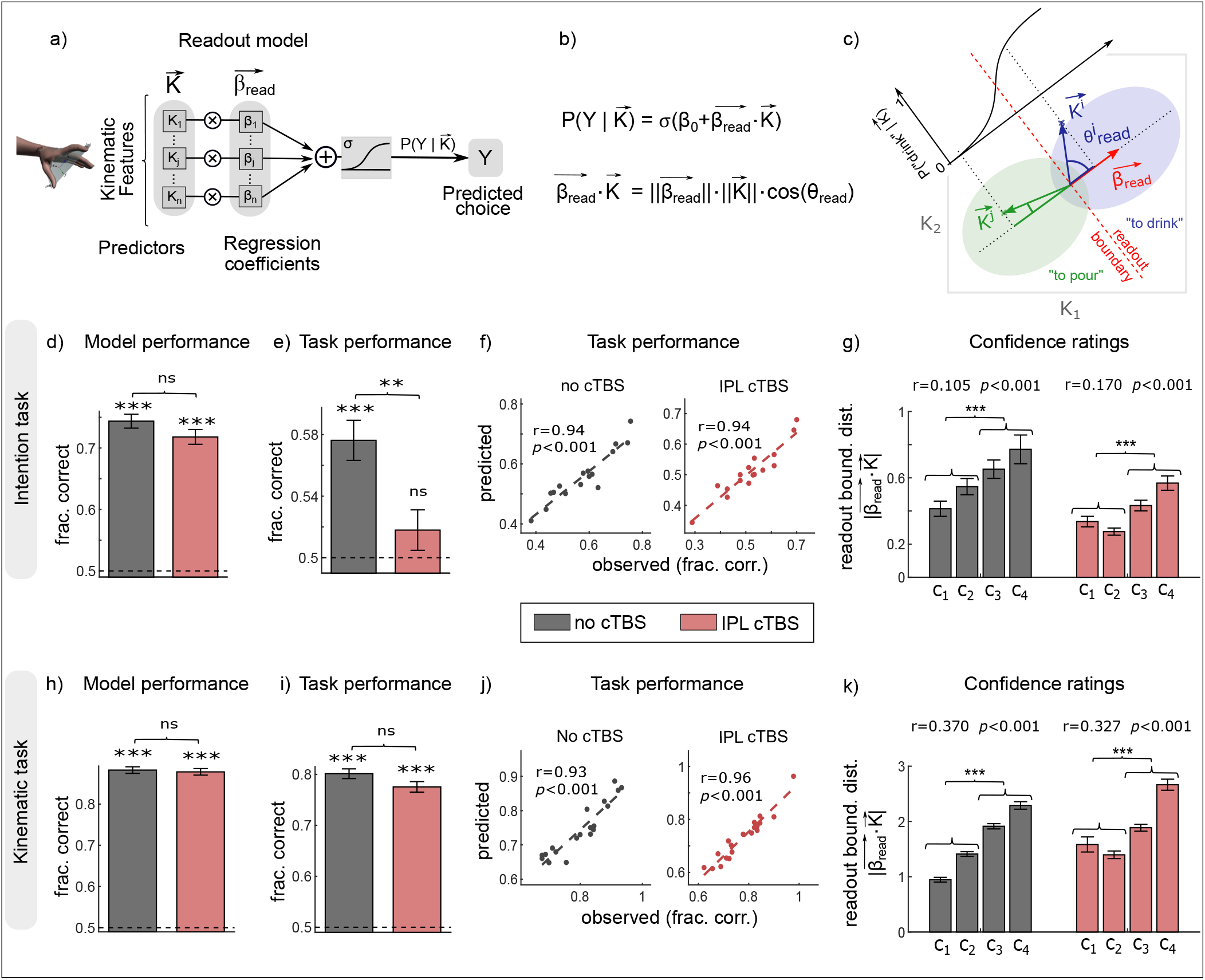
Readout model of intention discrimination from single-trial kinematics. **a-b** Readout model. Block diagram representation (a) and equation (b) of the logistic regression used to quantify intention information readout. **c** Sketch of the readout model in a simplified kinematic space spanning only two kinematic features. The two elliptic regions represent the intention choice conditional probability distributions of the two features. The readout vector 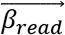 of regression coefficients indicates the direction in feature space that maximally discriminates observers’ choices. The direction and distance of the single-trial kinematic vector from the readout boundary determines, through the sigmoid logistic function, the probability of intention choice in that trial. **d** Performance of the readout model in predicting observers’ choices in the intention discrimination task quantified as fraction correct. **e** Task performance of participants quantified as fraction of correct intention choices. **f** Scatterplot of the relationship between the observed task performance and the one predicted by the readout model across individual participants in the intention discrimination task. **g** Distance of the single-trial kinematic vector from the readout boundary as a function of confidence ratings for the intention discrimination task. This distance was computed as module of the scalar product between single-trial kinematic vector and the readout vector. As shown in the panel c of Fig. 3, the larger this distance, the further from chance is the probability of intention choice predicted by the model. The green single-trial kinematic vector, for example, has a larger distance and thus a larger probability of intention choice than the blue kinematic vector. **h-k** Same as d-g for the kinematic discrimination task.

We fitted the readout model, separately for each subject, to single-trial intention choices and used confidence ratings reported by participants for independent validation of the model. Across trials and participants, model performance, measured as the fraction of intention choices correctly predicted by the model, was significantly above chance (Fig. 3d).

Although confidence ratings were not used for fitting model parameters, we also found a positive, trial-to-trial relationship between the observer’s confidence in their intention choice and the distance of single-trial kinematics vector from the readout boundary – the border that best separates kinematic patterns for the two intention choices (Fig. 3g). This suggests that intention choices on trials farther away from the readout boundary (and thus classified with greater confidence by the model) were also endorsed with higher confidence by human observers. Similar results were obtained for a readout model using single-trial differences in movement kinematics to predict kinematic discrimination performance (Fig. 3h-k).

To further test the predictive power of our readout model and evaluate how well it could account for task performance across sessions and tasks, we used the model to estimate the fraction of behaviorally correct trials. As the two alternatives were equiprobable and there was no response bias (see Supplementary Fig. 1), the fraction of behaviorally correct trials is *de facto* equivalent to the AUC^37,38^ (Fig. 3e, i). As shown in Fig. 3, we found close agreement between the observed and the predicted fraction of correct trials under no cTBS and IPL cTBS sessions in both the intention discrimination task (Fig. 3f) and the kinematic discrimination task (Fig. 3j).

Collectively, the analyses above suggest that our readout model was able to capture task performance, providing a plausible description of how well and how confidently observers performed the discrimination tasks.

### Transient disruption of IPL does not decrease sensitivity of intention readout to movement kinematics

Having verified that our readout model could account for intention discrimination performance, we next used it to adjudicate between alternative hypotheses regarding the functional consequences of IPL cTBS on action mindreading. The hypothesis of a decrease in sensitivity of intention readout to variations in movement kinematics following IPL cTBS predicts a weaker statistical dependency between single-trial kinematics and intention choices. We assessed this idea formally by comparing the fraction of intention choices correctly predicted by the model across sessions. The differences in the fraction of choices correctly predicted between no cTBS and IPL cTBS did not reach statistical significance (*p* = 0.07; Fig. 3d). These analyses suggest that the decrease observed in task performance following IPL cTBS cannot be accounted for by a decrease in sensitivity of intention readout to kinematics.

### Transient disruption of IPL causes misalignment of intention readout with respect to encoding

We next asked whether transient disruption of IPL alters ability to read the intentional significance of changes in movement kinematics. An intuitive visualization of how well readout captures intention-related information encoded in movement kinematics is provided by the angle between the encoding vector and the readout vector orthogonal to the readout boundary (Fig. 4a, b). The smaller the angle between these vectors, the larger the across-trial alignment between intention encoding and readout in kinematic space, and thus the larger the probability that intention-information is read out correctly. At the single-trial level, alignment can be computed as the angle between the single-trial kinematic vector and the readout vector. Given that the encoding vector expresses the direction around which intention information is expressed in kinematic space, increasing the angle between the encoding vector and the readout vector increases the angle between the single-trial kinematic vector and the readout vector (Fig. 4a, b). At the single-trial level, angles of 90° indicate that readout is totally unrelated to intention-information encoded in kinematics. Angles approaching 0° indicate full correct readout of intention information for ‘to drink’ trials (and totally incorrect readout of information for ‘to pour’ trials; Fig. 4a); angles approaching 180° indicate totally incorrect readout of intention-information for ‘to drink’ trials (and full correct readout of information for ‘to pour’ trials Fig. 4b). As shown in Fig. 4d, for no cTBS, single-trial angle distributions were centered 3° away from orthogonality, with ‘to pour’ and ‘to drink’ distributions only partly overlapping, and the majority of trials distributed in the correct readout angle range. For IPL cTBS, single-trial angles were centered only 1° away from orthogonality, with an almost complete overlap between intention-specific distributions and with about half of trials in the incorrect readout angle range (Fig. 4e). These data suggest that IPL cTBS impaired the ability to correctly readout intention information encoded in single-trial kinematics.

**Fig. 4.**
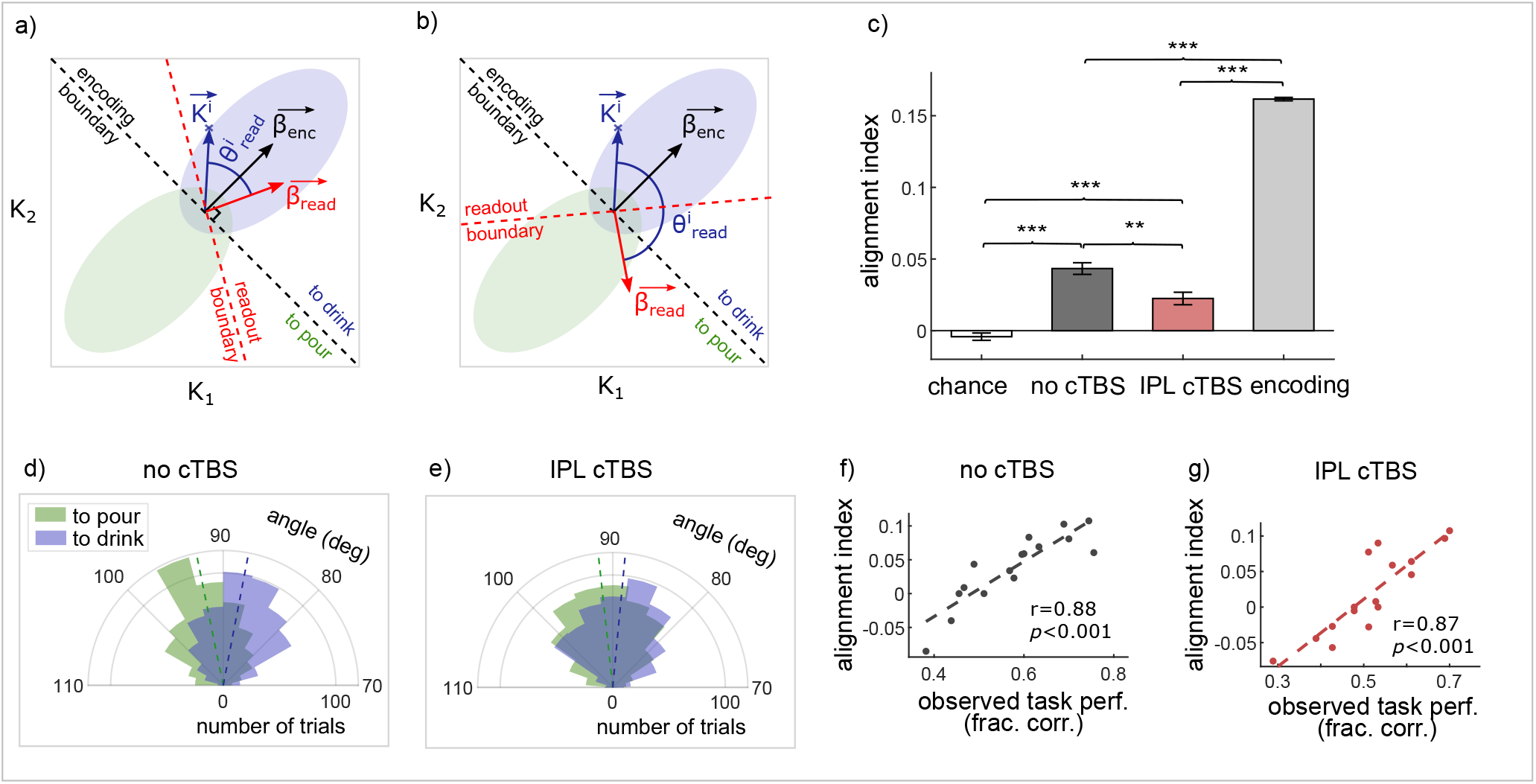
Misalignment of readout following IPL cTBS. **a-b** Diagrams illustrating the effect of misalignment in a simplified two-kinematic feature space. Conventions are as in Fig. 2e and 3c. The larger the angle between the encoding vector and the readout vector, the larger the angle between the single-trial kinematic vector and the readout vector. This justifies an alignment index based on the cosine of the angle between the single-trial kinematic vector and the readout vector, aligned in sign for each intention such that positive values correspond to correct readout of that intention. **c** Effect of IPL cTBS on the alignment index. For comparison, we show also the value of the alignment index for chance level (obtained after randomly permuting the intention choice labels) and the value of alignment index computed as the signed cosine of the encoding angle *θ_enc_*. As shown in Fig 2g, the angle *θ_enc_* between the encoding vector and the single-trial kinematic vector 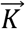 can be used to classify the intention of single-trial kinematics with 100% accuracy. Error bars indicate standard error of the mean. **d-e** Polar distribution of readout angles for ‘to pour’ and ‘to drink’ trials under no cTBS (d) and IPL cTBS (e). **f-g** Scatterplot of alignment indices against observed task performance across participants under no cTBS (f) and IPL cTBS (g).

To quantify these observations, we devised a single-trial alignment index based on the projection of the single-trial kinematic vector on the readout vector. First, we reflected the ‘to pour’ angle distribution across the 90° angle (so that, for example, a 95° angle was transformed into an 85° angle). Next, we pooled together ‘to drink’ and ‘to pour’ trials and computed the average cosine of the angle between the single-trial kinematic vector and the readout vector. With this formulation, positive alignment indices denoted correct readout and negative alignment indices denoted incorrect readout. Consistent with the finding that angle distribution was closer to orthogonality (90°) after IPL cTBS, results revealed a significant decrease of alignment after IPL cTBS (Fig. 4c). To rule out that such a decrease could be accounted for by differences in model performance, we repeated the analyses considering only those choices correctly predicted by the model. Results were qualitatively similar across sessions even when fully discounting the small difference in model performance between no cTBS and IPL cTBS (Supplementary Fig. 4g-i). For the kinematic discrimination task, no cTBS and IPL cTBS did not differ in alignment (Supplementary Fig. 4a-f). Together, these results suggest that transient disruption of IPL selectively misaligns intention readout with respect to encoding.

To substantiate the link between alignment and individual task performance, we quantified the fraction of behaviorally correct trials at the single-subject level as a function of alignment. Alignment was positively correlated with individual task performance (Fig. 4f, g).

The positive relationship between alignment and task performance at the single-subject level suggests that individual differences in task performance reflect differences in alignment between encoding and readout. However, individual differences could also reflect differences in the norm, or number, of non-zero readout coefficients. For example, poor performance may not only reflect poor alignment, but may result because non-zero readout coefficients are small in magnitude or number. To estimate the relative contributions of each of these factors to individual task performance, we performed a stepwise regression of task performance against the alignment index, the norm of the readout vector and the number of non-zero readout coefficients at the single-subject level. We found that alignment was the most important correlate of individual task performance; the explanatory power of the norm and the number of non-zero readout coefficients were two orders of magnitude lower (see Supplementary Data 5 and Supplementary Fig. 5).

Finally, we verified whether a decrease in alignment could predict a decrease in task performance after IPL cTBS at the individual subject level. Confirming this prediction, participants who experienced a larger decrease in alignment also experienced a larger decrease in task performance following IPL cTBS (Pearson correlation = 0.8, *p* < 0.001).

### Origins of misalignment between intention encoding and readout

To understand the origins of the decreased alignment induced by IPL cTBS, we further examined the distribution and concordance in sign of readout coefficients relative to encoding coefficients. We first assessed whether misalignment could result from a shift in the distribution of readout coefficients towards non-informative kinematic features, that is, whether a larger fraction of non-zero readout coefficients were assigned to non-informative individual kinematic features. The average fraction of non-zero readout coefficients assigned to informative kinematic features did not differ between no cTBS and IPL cTBS (Fig. 5a). Similarly, against the hypothesis of a shift of readout towards non-informative features, the norm of readout coefficients assigned to informative kinematic features was also similar across no cTBS and IPL cTBS (Fig. 5b). This suggests that transient disruption of activity in IPL did not alter the readout weight given to informative and non-informative kinematic features.

**Fig. 5.**
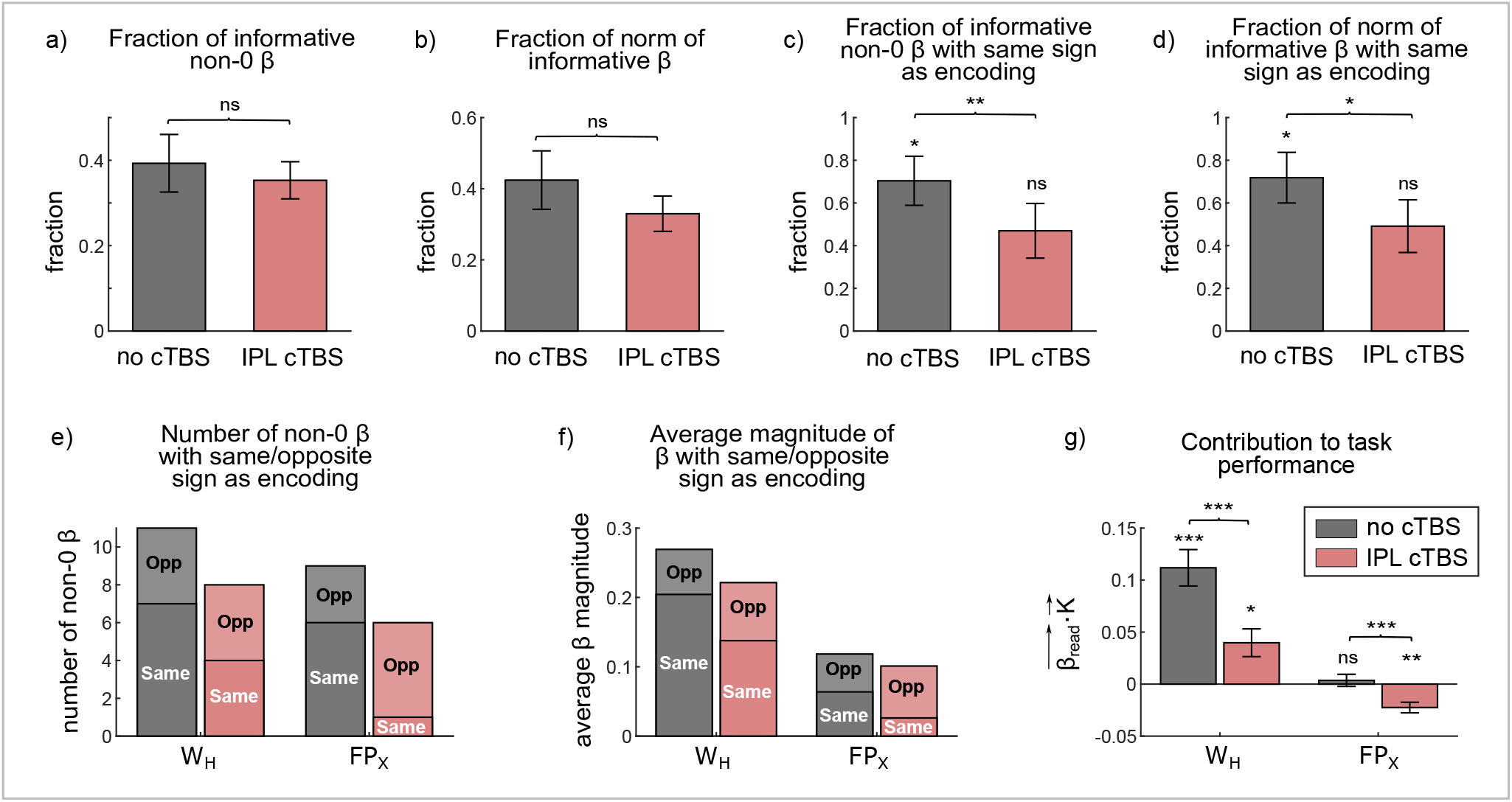
Origins of misalignment. **a-b** Fraction (a) and fraction of the norm (b) of non-zero readout coefficients assigned to informative features. **c-d** Fraction (c) and fraction of the norm (d) of non-zero readout coefficients assigned to informative features and correctly aligned with encoding. All fraction were computed on a subject basis and then averaged across subjects. **e-f** Readout weights with same/opposite sign with respect to encoding weights for the height of the wrist (WH) and the relative abduction/adduction of the thumb and the index finger, irrespective of wrist rotation (FPX). **g** Contribution of WH and FPX to intention task performance computed as the scalar product between the kinematic vector and the readout vector within that feature subspace. Error bars indicate standard error of the mean.

A second possible origin of decreased alignment is that IPL cTBS altered the mapping from informative kinematic features to intention choices. For example, a variation in a particular feature encoding ‘to drink’ (e.g., higher wrist height at 75% of movement duration; Fig. 2a), correctly read as ‘to drink’ before cTBS, might be incorrectly read as ‘to pour’ following cTBS. Geometrically, this would correspond to the readout vector and the encoding vector pointing in opposite directions (Fig. 4b), that is, the coefficients of the readout vector and of the encoding vector for the considered feature having opposite signs. Consistent with this hypothesis, the fraction of non-zero readout coefficients assigned to informative kinematic features with same sign was higher than chance in the cTBS condition and decreased to chance level following IPL cTBS (Fig. 5c). A similar pattern was found when considering the norm of readout coefficients assigned to informative kinematic features with same sign (Fig. 5d). In contrast, no differences between no cTBS and IPL cTBS were observed for the kinematic discrimination task (see Supplementary Fig. 6a-d). Together, these analyses indicate that cTBS to the left IPL altered the mapping of informative kinematic features to intention choices.

### Focus on specific kinematic features

To gain further insight into the inter-individual reproducibility of readout, we explored how specific features were read by different observers by computing cross-correlations between the readout weights of different participants. Observers showed moderately low cross-correlations between readout weights overall (see Supplementary Data 3), attesting to a diverse range of individual readout patterns. Interestingly, under no cTBS, cross-correlation values were significantly larger for informative features than for non-informative features. This indicates that readouts of non-informative features were more variable than those of informative features. After disruption of IPL, cross-correlation values no longer differed between informative and non-informative features, implying that readout variability was no longer reduced for informative features (see Supplementary Data 3).

Strikingly, under no cTBS, the two variables that were read out more consistently across observers (see Supplementary Data 3 and Supplementary Fig. 3c) were also more informative over a wider time range in terms of encoding (Supplementary Fig. 3a): the height of the wrist (WH) and the relative abduction/adduction of the thumb and the index finger, irrespective of wrist rotation (FPX). Under no cTBS, for WH, readout weights with the same sign with respect to encoding outweighed readout weights with opposite sign both in number and magnitude (Fig. 5e, f). This indicates that most WH readouts were appropriate. For FPX, in contrast, same sign readout weights were greater than those with opposite sign in number but not in magnitude (Fig. 5e, f), suggesting less appropriate readouts. IPL cTBS decreased the number and the magnitude of readout weights with the same sign for both variables. For FPX, but not for WH, this decrease was accompanied by an increase in number and magnitude of readout weights with the opposite sign relative to encoding.

To estimate the implications of these readout patterns for task performance, we computed the scalar product between the kinematic vector and the readout vector within the kinematic subspace formed by WH and FPX, respectively. The scalar product was assigned a positive value if the readout vector pointed toward the intention encoded in the kinematic vector and a negative value if the readout vector pointed in the opposite direction. Therefore, positive values of this index indicate a contribution of the considered kinematic variable toward the correct readout of intention information and thus correct choice. Negative values of this index indicate a contribution toward the incorrect read out of intention information and thus incorrect choice. As shown in Fig. 5g, whereas the contribution of WH to single-trial task performance was positive under no cTBS and remained positive after IPL cTBS, the contribution of FPX changed from null to negative, suggesting that, following IPL cTBS, incorrect readout of FPX contributed toward decreased task performance. For the kinematic discrimination task, IPL cTBS had no influence on how the most informative variables, including WH, were read out (Supplementary Fig. 6e-g). These results fit well with the above reported result of a selective decrease in alignment between intention encoding and readout after cTBS to the left IPL, and demonstrate how such decrease directly affected the most informative and most readout kinematic variables in the intention discrimination task.

## Discussion

While theoretical and experimental work suggest that action mindreading is mediated by the frontal-parietal action observation network, the specific neural computations involved in this ability have remained largely unknown and causally untested. Here we developed a novel approach, combining offline cTBS with analytical methods inspired by neural information coding, to reveal single-trial computations in left IPL and left IFG.

We found that cTBS applied over the left IPL, but not over the left IFG, impaired the ability to discriminate intention from variations in reach-to-grasp kinematics. Confirming the selectivity of the left IPL for intention discrimination, IPL cTBS did not affect the ability to discriminate kinematic variations unrelated to intention. These findings provide causal evidence of a functionally selective role of the left IPL in action mindreading. But what are the computations involved in action mindreading? What (and how) does the left IPL compute?

Our combined experimental and single-trial modeling results indicate that intention information is encoded in subtle, but highly informative variations in kinematic features. Under no cTBS, intention choice relied on the selective readout of such features at the single-trial level. The pattern of readout was sparse and idiosyncratic in that individual observers read different features. Across trials, however, informative features were read more, assigned larger readout weights and read more correctly than non-informative features. Notably, the two features more consistently read out across observers were also the two features informative over a wider time range in terms of encoding. Together, these findings suggest that knowledge about the natural statistics of intention coding – how intentions are encoded in movement kinematics – shapes intention readout, leading observers to place greater weight on features that carry more intention information.

Transient disruption of IPL did not impair ability to ‘see’ changes in movement kinematics, nor did it alter the relative weight given to informative versus non-informative features. Rather, it selectively decreased alignment between intention encoding and readout, affecting the observer’s ability to link variation in informative features to the correct intention. These results support a model of action mindreading in which the selection of the most informative kinematic feature occurs outside of the left IPL and in which the left IPL is selectively responsible for the correct readout of such features.

In contrast, IFG cTBS did not appear to have any measurable influence on intention readout. Both monkey and human studies relate left IFG to intention coding^6–8,16^. At least in humans, however, the left IFG seems to be less sensitive to intention readout compared to the left IPL^16^. Taken together with these previous reports, the lack of behavioral modulation to IFG cTBS in our study may indicate that, while potentially accessible to a classifier, intention-information in the left IFG is not causally related to task performance. Alternatively, it is possible that other brain regions can compensate for perturbation of the left IFG, possibly suggesting that computations performed in the left IFG, while functionally relevant for behavior, are not restricted to this area. To distinguish between these hypotheses, one could combine our readout model with dual-coil TMS and fMRI-TMS approaches.

The set of analytical methods developed in the current framework could be further generalized to examine how humans come to read a variety of mental states encoded in the movements of the eyes, mouth, hands and body. Moreover, our approach could be useful for developing intuition about how atypical encoding and readout link to deficits in social cognition^39^. For example, individuals with autism have difficulties perceiving, predicting and interpreting the actions of others (e.g., ^40^). The analysis and methods presented here could provide a useful tool for generating and testing alternative hypotheses about how altered readout computations affect ability to make inferences about others’ mental states.

## Methods

### Participants

Based on Cavallo et al.^41^, we decided *a priori* to collect data from at least 15 participants in each task. To achieve this, we had 20 participants perform the intention task and 20 participants perform the kinematic discrimination task. Three participants were removed from the sample as they did not complete all the three sessions. Additionally, two participants were unable to complete the cTBS sessions due to a too-high resting motor threshold (above 80% of maximal stimulator output). Thus, *n* = 16 for the intention discrimination task (10 females, 6 males, mean age 23, range 19-27 years) and *n* = 19 for the kinematic discrimination task (9 females, 10 males, mean age 24, range 20-28 years). All participants were right-handed according to the Edinburgh Handedness Inventory^42^ and had normal or corrected to normal vision. None of the participants reported neurological, psychiatric, or other medical problems or any contraindication to MRI or TMS^43,44^. Informed written consent was obtained in accordance with the principles of the revised Helsinki Declaration (World Medical Association General Assembly, 2008) and with procedures cleared by the local ethics committee (Comitato di Bioetica di Ateneo, University of Turin). All participants received monetary compensation for their time.

### Experimental design and procedures

The design of the intention discrimination and kinematic discrimination tasks was between-subjects, while effects of cTBS used a within-subject design. Participants assigned to each task underwent a high-resolution MRI structural scan, after which they attended three experimental sessions: no cTBS, cTBS to the left IPL and cTBS to the left IFG. During each of these sessions, participants completed the intention discrimination task (or the kinematic discrimination task, depending on task assignment) followed by a control contrast discrimination task. Participants sat in front of a 24-in. inch computer screen (resolution 1280 x 800 pixels, refresh frequency 60 Hz) at a distance of 50 cm in a dimly lit room. Each session lasted approximately 90 minutes and occurred at the same time of the day (±1 h) for each participant. Participant sessions were separated by one week, and session type order was randomized across participants.

#### MRI acquisition

T1-weighted scans were acquired using a 1.5 Tesla INTERA^™^ scanner (Philips Medical Systems) equipped with a 32-channel SENSE high-field head coil. Each high-resolution structural scan included 160 axial slices with an in-plane field of view (FOV) of 256 x 240 and a gap of 0 mm for a resolution of 1 x 1 x 1 mm (TR = 8.2 ms, TE = 3.80 ms, flip angle = 8 degrees). T1-weighted scans were used for the MRI-guided neuronavigation used to target cTBS stimulation sites (see below).

#### MRI-guided cTBS protocol

MRI-guided cTBS was administered using a 70-mm figure-eight coil connected to a Magstim Rapid2 stimulator (Magstim, Dyfed, UK). A SofTaxic NeuroNavigator system (EMS, Bologna, Italy) was employed to determine the coil position for all the ‘to-be-stimulated’ brain regions. Specifically, individual MRI scans were used to first construct scalp surface and skull landmarks of the left periauricular (A1), right periauricular (A2) and nasion (N) on the participant’s T1 MRI image. The brain scan was then normalized to Talaraich space and neuronavigation data were co-registered to measurements taken from the same points of reference (A1, A2, N) sampled from the participant’s scalp. The intensity for the cTBS protocol was set at 90% of the resting Motor Threshold (rMT), defined as minimal stimulation intensity producing motor evoked potentials (MEPs) of a minimum amplitude of 50 μV in the first dorsal interosseous (FDI) muscle^43^. To determine the rMT, for each participant for each stimulation session, we applied single pulse TMS over the left Primary Motor Cortex (M1) and recorded the MEPs from the right FDI muscle using a Biopac MP-150 (Biopac Systems, Inc., Santa Barbara, CA) through pairs of Ag–AgCl surface electrodes in a belly tendon montage. The rMT was determined by means of adaptive parameter estimation by sequential testing procedure (PEST) with the Motor Threshold Assessment Tool 2.0^45^. The coordinates for targeting left M1 (tal x = −44, y = −19, z = 53) were extracted from the neurosynth reverse inference map for the term ‘index finger’^46^. Following the rMT estimation procedure, cTBS was delivered to the left IFG and the left IPL. cTBS consisted of three pulses at 50 Hz repeatedly applied at intervals of 200 ms (5 Hz) for 40 s^33^. The two target sites of cTBS, left IFG and left IPL, were localized using neuronavigation based on MVPA classification results from Koul et al.^16^. To position the TMS-coil on the scalp location overlapping the target sites, the T1-weighted MRI-scan of each participant was loaded into SofTaxic Optic software (E.M.S. srl, Bologna, Italy). The stimulation sites were localized around the Talaraich stereotactic coordinates x = −46, y = 30, z = −11 for left IFG and x = −57, y = −30, z = 39 for left IPL (See Supplementary Fig. 7). In the IPL cTBS session and the IFG cTBS session, discrimination tasks were administered 5 min post cTBS, that is, in the time window in which maximal inhibitory effects of stimulation have been reported^33,47–50^.

#### Stimuli

Stimuli were selected from a dataset of 512 grasping acts obtained by recording 17 naïve participants reaching toward and grasping a bottle with the intent to pour some water into a small glass or to drink water from the bottle. Detailed apparatus and procedures are described in Cavallo et al.^41^. Briefly, reach-to-grasp movements were tracked using a near-infrared camera motion capture system with nine cameras (frame rate, 100 Hz; Vicon System) and concurrently filmed using a digital video camera (Sony Handy Cam 3-D, 25 frames/sec). Sixteen kinematic variables of interest were computed throughout the reach-to-grasp phase of the movement, from reach onset to reach offset^51^. A list of variables and how they are computed is reported in Supplementary Table 1 and in Supplementary Methods 1. Sixty grasping acts (grasp-to-pour, *N* = 30; grasp-to-drink, *N* = 30) were selected to satisfy the following requirements: i) within-intention distance was minimized (using the metric reported in Cavallo et al.^41^); ii) median split based on maximum wrist height led to a significant difference between “higher” and “lower” wrist height grasps (t_58_ = 11.2; *p* < 0.001); iii) maximum wrist height did not differ between intentions (*p* = 0.27). The corresponding movies, filmed from a lateral viewpoint, were used as stimuli in the intention discrimination task and in the kinematic discrimination task. Movies were edited with Adobe Premiere Pro CS6 (mp4 format, disabled audio, 25 frames per second, resolution 1,280 × 800 pixels) so that each movie clip started with the reach onset and ended at contact time between the hand and the bottle. Movement duration (mean ± sem = 1.04 ± 0.017 s, range = 0.84 to 1.36 s) did not differ between intentions (t_58_ = −0.30; *p* = 0.76).

#### Intention discrimination task

The intention discrimination task consisted of two blocks of 60 trials. Task structure conformed to a 2AFC design. Each trial displayed two reach-to-grasp acts in two consecutive temporal intervals: one interval contained a grasp-to-pour act, the other a grasp-to-drink act. Depending on block, participants had to indicate the interval (first or second) containing the grasp-to-drink or grasp-to-pour act. Each trial started with the presentation of a white central fixation cross for 1500 ms. Then, the first grasping act was presented followed by an inter-stimulus interval of 500 ms, after which the second grasping act was presented. After the end of the second video, the screen prompted participants to indicate the interval (first or second) containing the grasp-to-drink (or grasp-to-pour, depending on block) action by pressing a key. The prompt screen was displayed until response or for a maximum duration of 3000 ms. After response, participants were requested to rate the confidence of their choice on a four-level scale by pressing a key. Pairing of videos was randomized across trials and participants. To ensure that grasping actions could be temporally attended (i.e., to allow participants enough time to focus on movement start), 9, 11, or 13 static frames were randomly added at the beginning of each video. In order to equate video durations, static frames were also added at the end of each videos in a compensatory manner. Participants began the session by performing a practice block before the main experimental task. The order of the presentation of the blocks was counterbalanced across participants. Stimulus presentation, timing and randomization was controlled using E-prime V2.0 software (Psychology Software Tools, Pittsburgh, PA).

#### Kinematic discrimination task

The kinematic discrimination task included the same stimuli and design as the intention discrimination task, except that participant were asked to indicate the interval containing the grasp with higher (or lower, depending on block) peak vertical height of the wrist.

#### Control contrast discrimination task

To control for cTBS effects unrelated to action observation, such as integration of evidence favoring one alternative over time, participants performed a contrast discrimination task at the end of each session. The contrast discrimination task consisted of three blocks of 32 trials. Each trial started with the presentation of a fixation cross (1000 ms), after which two grey rectangles were displayed for 1000 ms on two consecutive intervals separated by a 500 ms inter-stimulus interval. In half of the trials, the rectangles had the same contrast (rgb = 100,100,100). In the other half of the trials, the difference in contrast was of 10, 15, 20 or 25 in the rgb space. For each trial, the participant had to indicate whether the contrast of the rectangles was ‘same’ or ‘different’ (within a 3000 ms window) and rate the confidence of their choice on a four-level scale by pressing a key. Results are reported in Supplementary Data 7.

### Data Analyses

#### Data preprocessing

Trials for which subjects failed to provide a response within 3000 ms were discarded from the analyses (0.5% of trials for the intention discrimination task and 0.1% of trials for the kinematic discrimination task). The first 25% of trials in each block were discarded prior to data analysis to allow time for all participants to become familiar with task and response mapping.

#### Computation of the ROC to quantify behavioral performance

To quantify each participant’s behavioral performance in the discrimination tasks, we first combined participants’ responses and confidence ratings to estimate points on an empirical ROC curve^37,38^. The decision variable for the ROC determination was computed by combining the binary discrimination response (reporting of first/second interval) and the four-level confidence rating into a single eight-level rating response that was used as the decision variable for the ROC calculation. This eight-level rating response associates levels 1-4 (from low to high confidence) to discrimination responses reporting “first interval”, and levels 5-8 to discrimination responses reporting “second interval” (from low to high confidence). We then estimated the performance of each participant by computing the AUC using Matlab’s function *perfcurve*.

#### Single-trial kinematic vector

To model single-trial kinematics, we first averaged, for each grasping act, the 16 kinematic variables of interest over four intervals of 25% of the normalized movement time (defined from reach onset to reach offset). Next, for each trial, we combined the kinematic features associated with the two grasping acts in a 64-dimensional kinematic vector, 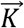, defined as:

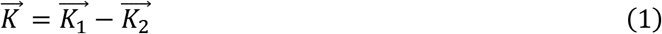

where 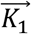 and 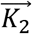 denote the vectors of kinematic features associated to the first and second reach-to-grasp act displayed in the trials. This definition, used in all of our logistic regression analyses, reflects the assumption that, in a 2AFC task, choices are based on comparative judgements.

Using more detailed regression models that employed the kinematic features of the two grasping acts did not improved model predictability (see Supplementary Data 2).

#### Logistic regression models

We analyzed encoding and readout using two sets of logistic regression models: encoding and readout models. Logistic regression^36^ is a linear regression for log-odds, and is a standard probabilistic approach to classification. Logistic regression models are powerful for explaining behavioral strategies^52^. They confer several advantages in modeling the dependence of a random binary variable, such as observers’ choice, on one or more explanatory variables^53^. For example, they assume binomial noise – the most natural noise model for binary responses; they combines predictor variables linearly; they can be robustly fit to data; they have a graded nonlinearity, which allows for a modulation of probabilities different from an all-or-none binarization. The latter property was particularly suitable to the readout model because in our data discrimination performance was positively correlated with confidence ratings in the no cTBS session (Spearman’s correlation: *p* < 0.05 and *p* < 0.001 for the intention and kinematic discrimination tasks, respectively), suggesting a graded nature of the response probability as a function of the kinematic evidence. To aid comparison between encoding and readout models, we also used logistic regression for modeling encoding. Versions of the encoding models based on other formulations, such as linear discriminant analyses, were also built and tested, and yielded qualitatively similar results.

The logistic model expressed the probability of a binary stochastic variable *Y*, where *Y* takes the values ‘to drink’ and ‘to pour’ for intention discrimination, as a sigmoid transformation of the sum of the components of the single-trial kinematic vector 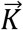. The equation of the model was as follows:

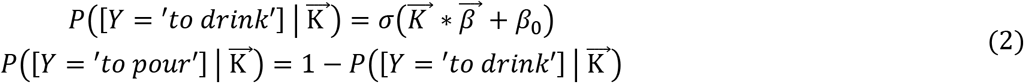

where *σ* is the sigmoid function, 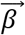 is the vector containing the values of the regression coefficients of each kinematic feature, and *β_0_* is the bias, kinematic independent, term.

#### Training logistic regression models

Training and evaluation were performed similarly for both sets of models and for both discrimination tasks. Each model was trained on the set of 90 trials retained for analyses. We z-scored the single-trial kinematic vectors within each model in order to avoid penalizing predictors with larger ranges of values. To avoid over-fitting, we trained each model using elastic-net regularization, with a value of α = 0.95 for the elastic net parameter, which provides sparser solutions in parameter space^54^. We verified that our results were robust to the choice of elastic net regularization (see Supplementary Data 3). The free parameter *λ*, which controls the strength of the regularization term, was estimated for each model using leave-one-out cross-validation. We retained for each model the value *λ*_min_ associated to the minimum mean cross-validated error. Models were then trained on all 90 trials with the retained regularization term. Logistic regression was implemented using *R glmnet* package^55^. In the main text, we report the results obtained by applying this training procedure as it gives only one set of regression coefficients per analyzed case and it is therefore easier to interpret. However, qualitatively similar results were obtained when using leave-one out cross validation on the entire procedure (on top of the cross validation used for the determination of the λ parameter; see Supplementary Data 3).

The following sections describe encoding and readout models with reference to the intention discrimination task. The procedures for the kinematic discrimination task were identical.

#### Encoding model

The encoding model expressed the probability of the grasping act displayed in the first interval of a given trial being ‘to drink’ as a function of the kinematic vector measured in the same trial. Having verified that intention information slightly varied as a function of video pairings (which was randomized across trials and participants), we trained the encoding model separately on each set of video pairings presented in each session to each observer. We used the encoding model to evaluate the overall amount of intention information in movement kinematics (Fig. 2g, 4c).

#### Readout model

The readout model expressed the probability of intention choice in a given trial as a function of the kinematic vector measured in the same trial. We trained the readout model separately for each observer in each session. To model intention choice as a function of single-trial kinematics, we trained the readout model using all 64 kinematic features (16 kinematic variables at four time points).

#### Evaluation of model performance

To quantify model performance (Fig. 2g, 3d), we computed for each trial the most likely value of the variable *Y* by taking the argmax over *Y* of 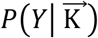 in Eq. 2. This parameter provides an estimate of the model prediction (prediction of the actual intention for the encoding model; prediction of observer’s choice for the readout model) on a single trial. We then quantified model performance as the fraction of correct predictions computed over all the trials.

#### Computation of the statistical significance of model performance

To assess the statistical significance of model performance, we estimated model performance under the null-hypothesis that trial labels (actual intention for the encoding model; observer’s choice for the readout model) can be permuted across trials without affecting the model performance. For each model, we performed 100 random permutations and fitted logistic regression to these randomly permuted data sets. This provided a null-hypothesis distribution of model performance values that was then used to compute non-parametric *p*-values of the model performance on the true data. To further check that that the regularization was working well and that the regression coefficients with non-zero value were meaningful^36^, we took the absolute value of each individual regression coefficient obtained in the permuted dataset to build a distribution of absolute values of regression coefficient expected under the null-hypothesis of no relationship between the kinematics and the variable *Y*. We verified that all non-zero beta coefficients had an absolute value that exceeded the 95th percentile of this null-hypothesis distribution.

#### Computation of task performance predicted by the readout model

In Fig. 3f, j, we used the readout model to estimate, for each participant, the fraction of behaviorally correct trials. Using the logistic readout model (Eq. 2), we computed for each trial the probability of each intention choice. Then we averaged across all trials and participants the probability of the correct intention choice.

#### Classification of individual kinematic features as informative

To evaluate the informativeness of individual features about intention (which is used for Fig. 5a-g, Supplementary Fig. 3a, b, and Supplementary Fig. 6), we used a single-feature encoding model implemented using Matlab’s *glmfit* function. Each model was trained on the full set of 90 trials retained for analyses. We z-scored the single-trial kinematic vectors. Significance of the regression coefficient was assessed with t-statistics. We retained as informative kinematic features whose regression coefficients were found to be significant (*p* < 0.05) in all video pairings. In Supplementary Fig. 3a, b, variables are ranked in terms of their encoded information. Ranking was determined by computing the average magnitude of the regression coefficient of each informative feature across all video pairings and then summing the obtained magnitudes across features belonging to the same kinematic variable.

#### Contribution of individual kinematic variables to task performance

Fig. 5g and Supplementary Fig. 6g visualize the contribution of individual kinematic variables to task performance. Single feature contribution to task performance was computed as the scalar product between the kinematic vector and the readout vector calculated within the feature subspace formed by the features of the considered kinematic variable (e.g., W_H_ at 25%, 50%, 75% and 100% for W_H_). Positive values of this index imply a positive contribution of the variable towards enhancing task performance; negative values imply a negative contribution towards decreasing task performance.

#### Statistics

##### Subject level analyses of task performance (Fig. 1d, e)

We assessed the significance of task performance above chance level in each session with one-tailed one sample t-tests. We assessed the significance of the decrease in task performance induced by cTBS with one-tailed paired sample t-tests.

##### Trial level analyses-logistic model

Unless otherwise stated, differences in values across sessions were assessed using non-parametric permutation tests, based on constructing a null-hypothesis distribution of differences in values after randomly permuting trial labels across sessions.

Comparisons of task performance and model performance across sessions was performed by a permutation test, where a null-hypothesis distribution of values with no association between trials and sessions was constructed by randomly permuting the session labels across trials. We found empirically that different subject choices in different trials were statistically independent even when considering consecutive trials (mutual information test, p < 0.05). However, to conservatively incorporate any residual effects of correlations across almost consecutive trials in the null-hypothesis distribution, we permuted sessions labels within blocks of five consecutive trials. For all tests, the null-hypothesis distribution was computed using 10^4^ random permutations. We used permutation tests because they do not make assumptions about the nature of the distribution of the individual values and are therefore applicable to a wide range of quantities, from the distribution of regression weights to behavioral performance. We assessed the significance of differences across sessions using two-tailed statistics and the significance of increase or decreases of quantities using one-tailed statistics. All *p*-values of comparisons across sessions were Holm-Bonferroni corrected for two comparisons (no cTBS vs. IPL cTBS, and no cTBS vs. IFG cTBS).

##### Significance of correlations

Significance of Pearson correlation values and step-wise regression coefficients were assessed using two-tailed parametric Student statistics^56^ implemented in the MATLAB functions *corr* and *stepwisefit*, respectively. We assessed significance of Spearman correlations using the two-tailed permutation distribution^57^ implemented in the MATLAB function *corr*.

##### Conventions for plotting p values

In all figures, * indicates *p* < 0.05, ** indicates *p* < 0.01, and *** indicates *p* < 0.001

## Author Contributions

C. B. and S. P. conceived and supervised the project. C. B. and A. C. designed the experiments. S. P., C. B, J-F. P. and M. V. designed the analyses. A. K. and M. S. performed the experiments. J-F. P. performed the analyses, with contributions from M. V. and A. K. A. A. contributed methods. C. B., S. P. and J-F. P wrote the manuscript. All authors contributed to the content and writing of the Supplementary Information.

