## Supplementary Information for "Transient disruption of the inferior parietal lobule impairs action mindreading"

### *Supplementary Methods*

***Supplementary Methods 1.*** *Computation of kinematic variables*

We used a custom software (Matlab; MathWorks, Natick, MA) to compute two sets of parameters of interest: $F_{global}$ and $F_{local}$ parameters. $F_{global}$ parameters were expressed with respect to the global frame of reference, i.e., the frame of reference of the motion capture system. Within this frame of reference, we computed the following parameters:

a) Wrist Velocity, defined as the module of the velocity of the wrist marker (mm/sec);
b) Wrist Height, defined as the z-component of the wrist marker (mm);
c) Wrist Horizontal Trajectory, defined as the x-component of the wrist marker (mm);
d) Grip Aperture, defined as the distance between the marker placed on thumb tip and the one placed on the tip of the index finger (mm).

To provide a better characterization of the hand joint movements, the second set of parameters was expressed with respect to a local frame of reference centered on the hand (i.e., $F_{local}$). Within $F_{local}$ we computed the following parameters:

e) x-, y-, and z-thumb defined as x-, y- and z-coordinates for the thumb with respect to $F_{local}$ (mm);
f) x-, y-, and z-index defined as x-, y- and z-coordinates for the index with respect to $F_{local}$ (mm);
g) x-, y-, and z-finger plane defined as x-, y- and z-components of the thumb-index plane, i.e., the three-dimensional components of the vector that is orthogonal to the plane, providing information about the abduction/adduction movement of the thumb and index finger irrespective of the effects of wrist rotation and of finger flexion/extension;
h) x-, y-, and z-dorsum plane defined as x-, y- and z-components of the radius-phalanx plane, providing information about the abduction, adduction and rotation of the hand dorsum irrespective of the effects of wrist rotation.

***Supplementary Table 1.*** *Kinematics parameters of interest. Kinematic variables were computed throughout the reach-to-grasp phase of grasp-to-pour and grasp-to-drink acts, from reach onset to reach offset.*

| **Kinematics parameter** | **Definition** |
|---|---|
| Wrist velocity | module of the velocity of the wrist marker (mm/sec) |
| Wrist height | z (i.e. up-down) component of the wrist marker (mm) |
| Wrist horizontal trajectory | x (i.e. left-right) component of the wrist marker (mm) |
| Grip aperture | distance between the marker placed on the tip of the thumb and the marker placed on the tip of the index finger (mm) |
| x-, y-, z-index | x-, y- and z-coordinates for the index (mm) |
| x-, y-, z-thumb | x-, y- and z-coordinates for the thumb (mm) |
| x-, y-, z-finger plane | x-, y- and z-components of the thumb-index plane (mm) |
| x-, y-, z-dorsum plane | x-, y- and z-components of the radius-phalanx plane (mm) |

### *Supplementary Data*

***Supplementary Data 1.*** *Analysis of the effect of cTBS stimulation on kinematic discrimination performance at different levels of task difficulty*

To rule out that cTBS effects in the kinematic discrimination task were limited by the overall high task performance, we repeated our analysis subsampling trials at different levels of difficulties. Using methods defined in Section "Encoding of intention-related information" of the main text, we computed the distance of each trial from the encoding boundary and ordered trials in order of increasing distance, from the nearest (and thus less discriminable) to the farthest (and thus most discriminable). Next, we split trials in five levels of difficulty and computed AUC values (and percent correct) for each quintile in each session. We found no significant effects of cTBS across levels and sessions ($p > 0.1$). This suggests that the lack cTBS effects in the kinematic discrimination task was not related to task difficulty.

***Supplementary Data 2.*** *Systematic comparison of readout models that use different numbers of kinematic features in different ways*

In all logistic regression analyses presented in the main text, the single trial kinematic vector was constructed as the differences between the kinematic features of the reach-to-grasp acts displayed in the first and second interval of each trial ($\overrightarrow{K} = \overrightarrow{K_1} - \overrightarrow{K_2}$). We assessed whether a more complex model, using the full set of kinematic variables of the first and second grasping act independently (that is, $\overrightarrow{K} = [\overrightarrow{K_1}, \overrightarrow{K_2}]$), rather than their difference, would achieve better performance. The same regularization procedure, described in Methods in the main text, was applied. Performance of the model using the full set of kinematic features was not significantly different from that of the model using the difference in kinematics in any condition of the intention discrimination task ($p > 0.5$). For the kinematic discrimination task, the model using the full set of features showed a small advantage for no cTBS and IPL cTBS sessions (0.88 vs 0.91, $p < 0.001$ for no cTBS, 0.88 vs 0.90 $p < 0.001$ for IPL cTBS). Both approaches achieved similarly high correlations between predicted and observed task performance ($p < 0.001$ in all cases, Fig SI2). Given that the model using the full set of kinematic features had twice as many predictors as the model using the difference (128 vs 64 dimensions), but only led to a null to marginal increase in model performance, for the sake of parsimony, we used the simpler model in all analyses.

We also explored whether the number of time-bins used for the discretization of the temporal evolution of kinematic variables influenced model performance. We compared the performance of the four-time-bin readout model, used in all our analyses presented in the main text and supplemental figures, with that of models based on a more-detailed time discretization (six or eight time bins). Six- and eight-time bin models performed no better than the four-time bin model ($p > 0.05$).

***Supplementary Data 3.*** *Similarity of readout weights across observers in different conditions and with different model regularizations*

To gain further insight into the robustness and inter-individual reproducibility of the readout regression coefficients, we computed cross-correlations between the readout weights of different participants under different control analyses.

We first computed the cross-correlation between the readout weights for each different pairs of participants. We found that participants showed overall moderate to low cross-correlations between their readout weights under no cTBS in the intention discrimination task (mean $\pm$ sem across all pairs of subjects was $0.06 \pm 0.02$). To investigate whether the small correlation values were due to the regularization, we repeated the above analysis using different values of the hyper-parameter $\alpha$ (the one bridging continuously between Ridge regularization for $\alpha$ close to zero and Lasso regularization for $\alpha$ close to one) ranging from 0.5 to 1. This range of elastic net hyper-parameter $\alpha$ covers the range of regularization further away from the Ridge and closer to the Lasso one, which is the most suitable range given our number of trials and predictors. Cross-subject correlations slightly increased with lower $\alpha$ values, but remained always below 0.1. Similar results were obtained for IPL cTBS and IFG cTBS. As expected by design, cross-correlations in the kinematic discrimination task were much higher (mean $\pm$ sem across all pairs of subjects was 0.39 $\pm$ 0.02 in no cTBS session), and remained high for all $\alpha$ values. These analyses corroborate the idea that the small correlation values in the intention discrimination task reflected genuine cross-subject differences.

We also considered whether cross-correlation of readout patterns differed between non-informative and informative kinematic features in the intention discrimination task. We considered separately the 19 out of 64 features that carried significant intention-related information ($p < 0.05$) and computed the cross-subject correlation value in this subset of variables. Next, we compared this value with that obtained when considering the 19 non-informative features with the smallest encoding readout coefficients. Under no cTBS, cross-correlation values were significantly larger for informative features than for non-informative features ($p < 0.05$). After IPL cTBS, cross-correlation values did not differ ($p = 0.14$) between informative and non-informative features. In the kinematic discrimination task, cross-correlations values were significantly larger for informative features than for non-informative features in all conditions ($p < 0.001$).

To check whether the pattern of readout weights was robust to the choice of the elastic net hyper-parameter $\alpha$, we computed for each participant the correlation between the readout weights for $\alpha$=0.95 with the readout weights obtained with values of $\alpha$ ranging from 0.5 to 1. Correlation values decreased with $\alpha$ values but remained higher than 0.9 for all $\alpha > 0.5$. Altogether, these results indicate that readout results were robust to the choice of hyper-parameter $\alpha$.

***Supplementary Data 4.*** *Cross-validated model performance and cross-validated alignment results*

In the analyses reported in the main text, model performance was evaluated on the same set of trials used for training. To control for overfitting, we recomputed all our main analyses using a different partitioning of data into training and testing (leave-one-out cross-validation). The pattern of results obtained with this cross-validated approach was consistent with that reported in the main text. In particular, the cross-validated performance of encoding models remained close to 100% for both the intention and the kinematic discrimination tasks (99.5$\pm$0.1% and 98.1$\pm$0.2% for the intention discrimination task and kinematic discrimination task, respectively). The cross-validated performance of readout models also remained significantly above chance in all sessions in both tasks ($p < 0.001$, adjusted for the three comparisons in each task). The correlation between observed and predicted task performance also remained significant in all cases ($p < 0.001$). Moreover, alignment was still significantly decreased after IPL cTBS as compared to no cTBS ($p < 0.01$).

***Supplementary Data 5.*** *Alignment is the main predictor of task performance*

To establish whether alignment was a predictor of task performance, we computed, for each participant, the Pearson correlation between the fraction of behaviorally correct trials and the alignment index. Alignment indices for participants with zero readout vectors were set to zero as in this case no information can be read out. Results, plotted in Fig. 4f and Supplementary Fig. 5, showed that the alignment index was highly correlated with task performance (Pearson correlation = 0.89, $p < 0.001$ for the no cTBS condition). Model parameters related to the strength of readout, but not to alignment, namely, the norm of readout vector and the number of non-zero readout regression coefficients, were also considered (Supplementary Fig. 5). These parameters showed a much weaker, borderline significant, correlation with task performance under no cTBS and no correlation after IPL cTBS in the intention discrimination task. Comparable results were found in the kinematic discrimination task (Supplementary Fig. 5).

To rank the importance of the above considered model parameters in predicting changes in task performance across conditions, we conducted stepwise linear regression analyses of the log of the ratios between IPL cTBS and no cTBS task performance of each individual participant. In both the intention discrimination task and in the kinematic discrimination tasks, the alignment index was ranked as the most important predictor (Supplementary Table 2). Only in the kinematic discrimination task, the norm of the readout vector added a significant contribution as second predictor ($p = 0.003$). The number of non-zero readout regression coefficients never added predictive power ($p > 0.05$). Collectively, these analyses support the conclusion that the alignment index was the main predictor of the change in task performance between no cTBS and IPL cTBS for each individual participant.

***Supplementary Table 2.*** *Results of stepwise regression analyses of the log of the ratio between IPL cTBS and no cTBS task performance of each individual participant using the alignment index, the norm of the readout vector and the number of non-zero readout coefficients as predictors.* The table reports, for each predictor, the value of the stepwise regression coefficient, its standard error and its p value. The predictors are ordered and listed from top to bottom, in terms of importance as computed by the stepwise regression.

| **Intention discrimination** | **Predictor** | **Coefficient** | **Standard error** | **p- value** |
|---|---|---|---|---|
| | alignment | 4.031 | 0.811 | < 0.001 |
| | norm readout vector | -0.039 | 0.026 | 0.169 |
| | n. non-zero readout coefficients | -0.067 | 0.036 | 0.095 |

| **Kinematic discrimination** | **Predictor** | **Coefficient** | **Standard error** | **p- value** |
|---|---|---|---|---|
| | alignment | 3.811 | 0.676 | < 0.001 |
| | norm readout vector | -0.096 | 0.027 | 0.003 |
| | n. non-zero readout coefficients | -0.045 | 0.028 | 0.138 |

***Supplementary Data 6.*** *Results for IFG cTBS*

In both the intention discrimination task and in the kinematic discrimination task, logistic regression analyses showed similar results for IFG cTBS and no cTBS. In particular: i) readout model performance under IFG cTBS was significantly higher than chance ($p < 0.001$) and did not differ from model performance under no cTBS ($p > 0.05$); ii) the correlation between observed and predicted task performance was as high for IFG cTBS as for no cTBS (and IPL cTBS; see Supplementary Fig. 2); iii) confidence ratings and the distance of the single-trial kinematic vector from the readout boundary showed a positive trial-to-trial correlation under IFG cTBS (Spearman correlation; $p < 0.001$ for both the intention discrimination task and the kinematic discrimination tasks); iv) alignment of readout to encoding as measured by the alignment index was similar under IFG cTBS and no cTBS ($p > 0.05$).

***Supplementary Data 7.*** *Results of the contrast discrimination task*

Trials for which subjects failed to provide a response within 3000 ms were discarded from the analyses (0.5% of trials performed by subjects in the intention discrimination group and 0.3% of trials performed by subjects in the kinematic discrimination group). Furthermore, the first 25% of trials of each block were discarded, as for all analyses performed for the intention discrimination task and the kinematic discrimination task. Task performance, as determined by the AUC revealed no influence of IPL cTBS or IFG cTBS in either task ($p > 0.05$).

### Supplementary Figure Captions

***Supplementary Fig. 1.*** *Absence of response bias.* **a-b** Fraction of 'to drink' answers in the intention discrimination task (a) and 'higher' answers the kinematic discrimination task (a) in each experimental session. Results are reported as mean ± sem across subjects.

***Supplementary Fig. 2.*** *Comparison between models using the difference in kinematic features and models using all the kinematic features of the two reach-to-grasp acrs.* **a-b** Scatterplot of the relationship between the observed task performance and the one predicted by the readout model across individual participants in the intention discrimination task (a) and in the kinematic discrimination task (b).

***Supplementary Fig. 3.*** *Ranking of features with respect to encoding and readout.* **a-b** Sum of the absolute value of encoding regression coefficients over time bins encoding significant intention (a) and wrist height (b) discriminative information. Features are ordered left to right from most informative to least informative. **c-d** Number of observers who read out a specific variable over time bins in the intention discrimination task (c) and in the kinematic discrimination task (d). Features are ordered left to right from most to least readout.

***Supplementary Fig. 4.*** *Additional analyses on the effect of IPL cTBS on alignment.* **a-b** Polar distribution of readout angles for 'higher' and 'lower' trials under no cTBS (a) and IPL cTBS (b) conditions in the kinematic discrimination task. **c** Effect of IPL cTBS on alignment in the kinematic discrimination task. **d-e**. Polar distribution of readout angles for 'higher' and 'lower' trials under no cTBS (d) and IPL cTBS (e) conditions in the kinematic discrimination task considering only choices correctly predicted by the model. **f** Effect of IPL cTBS on alignment in the kinematic discrimination task considering only choices correctly predicted by the model. **g-h** Polar distribution of readout angles for 'to pour' and 'to drink' trials under no cTBS (g) and IPL cTBS (h) conditions in the intention discrimination task considering only choices correctly predicted by the model. **i** Effect of IPL cTBS on alignment in the intention discrimination task considering only choices correctly predicted by the model. For graphical representation, in panels a-b, d-e, g-h, the 70-110° angle range of polar distributions are expanded to a semi-circle.

***Supplementary Fig. 5.*** *Correlation of model parameters with task performance.* ***a-b*** Scatterplot of the alignment index, the norm of readout vector and the number of non-zero readout regression coefficients against observed task performance across participants under no cTBS and IPL cTBS conditions in the intention discrimination task (a) and in the kinematic discrimination task (b).

***Supplementary Fig. 6.*** *Quantification of alignment in the kinematic discrimination task.* **a-b** Fraction (a) and fraction of the norm (b) of non-zero readout coefficients assigned to informative features. **c-d** Fraction (c) and fraction of the norm (d) of non-zero readout coefficients assigned to informative features and correctly aligned with encoding. All fraction were computed on a subject basis and then averaged across subjects. **e-f** Readout weights with same/opposite sign with respect to encoding for the wrist height ($W_H$) and the wrist horizontal trajectory ($W_{HT}$). **g** Contribution of $W_H$ and $W_{HT}$ to task performance in kinematic discrimination computed as the scalar product between the kinematic vector and the readout vector within the kinematic subspace of the variable. Error bars indicate standard error of the mean.

***Supplementary Fig. 7.*** *Target sites of cTBS.* Individual T1-weighted scans used for MRI-guided neuronavigation in the intention discrimination task and in the kinematic discrimination task. Target sites for the left IFG and the left IPL are marked by red crosses.

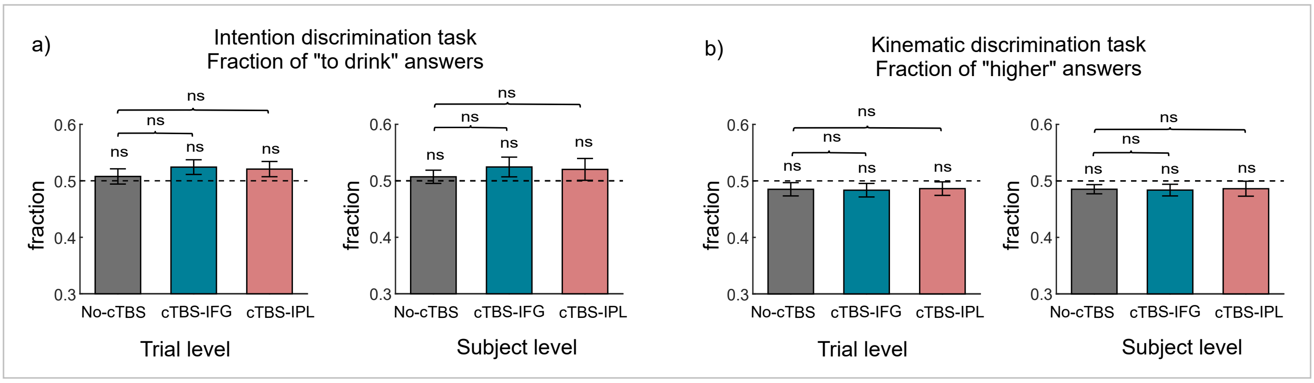

Supplementary Figure 1

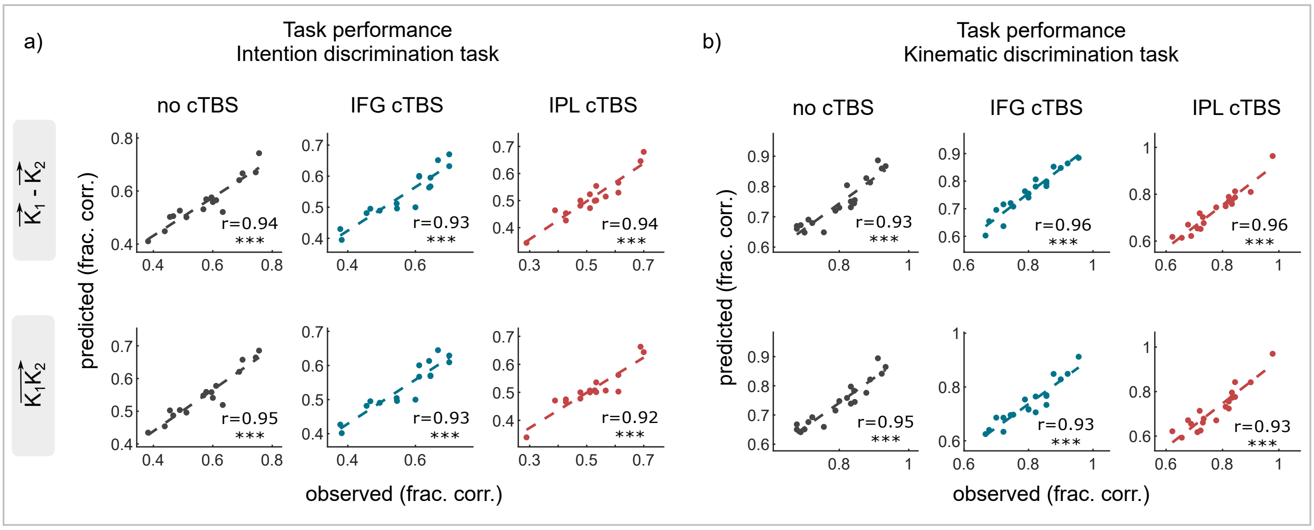


Supplementary Figure 2

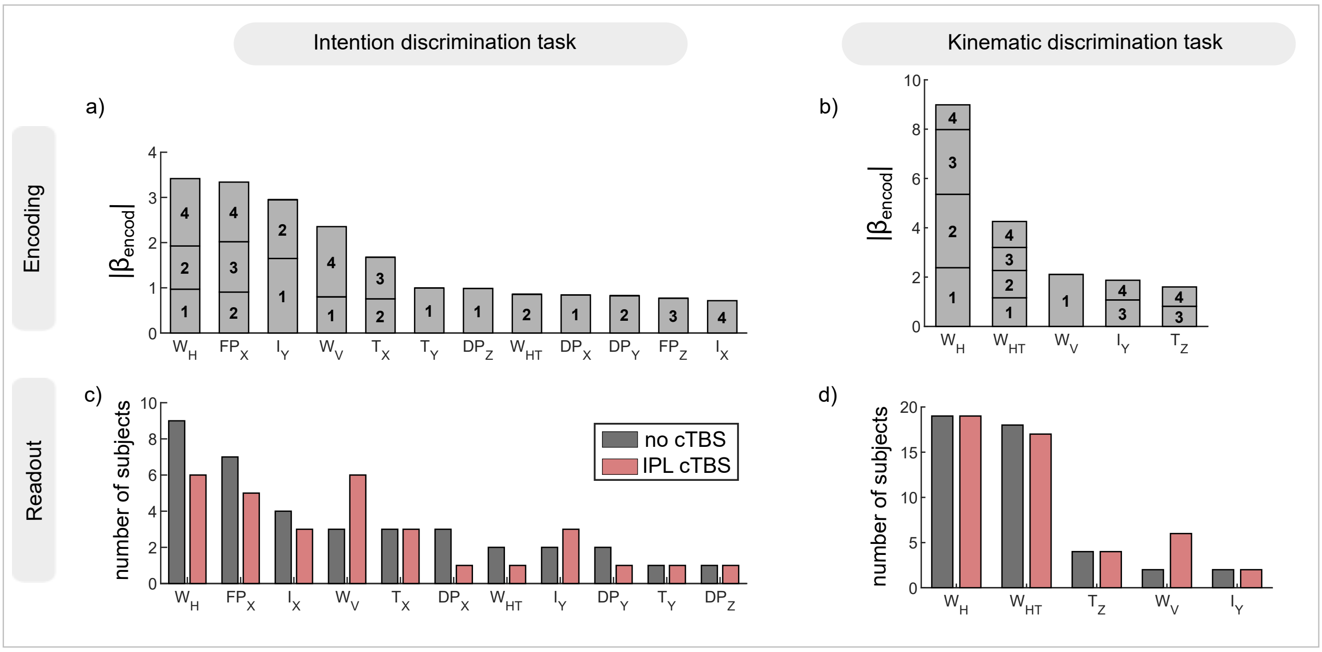


Supplementary Figure 3

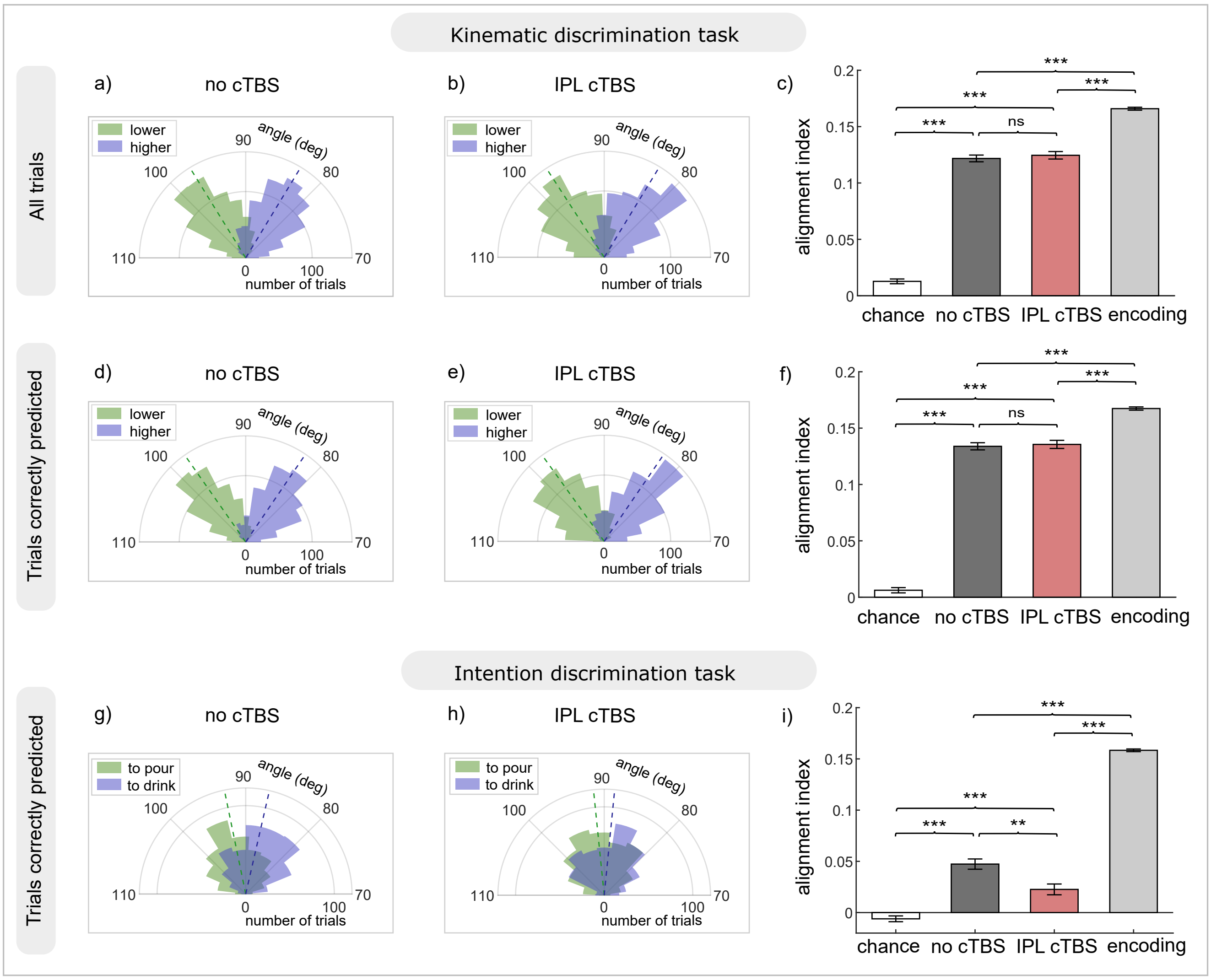


Supplementary Figure 4

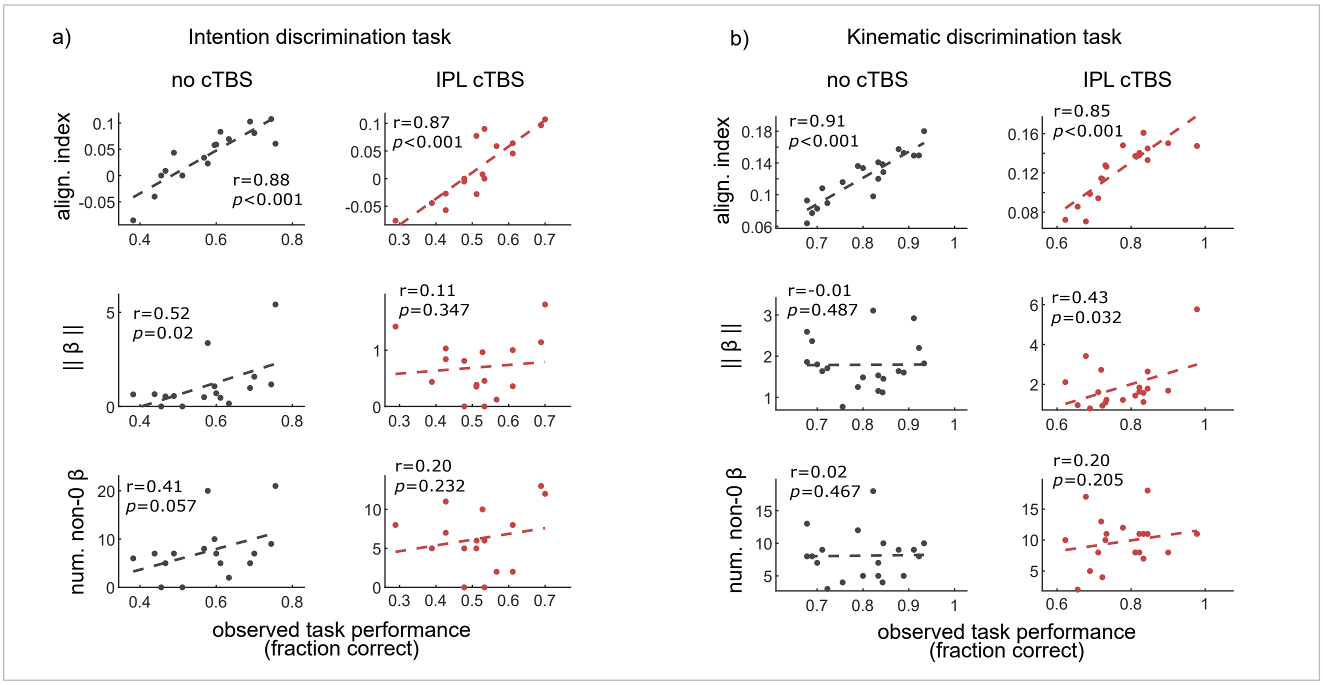


Supplementary Figure 5

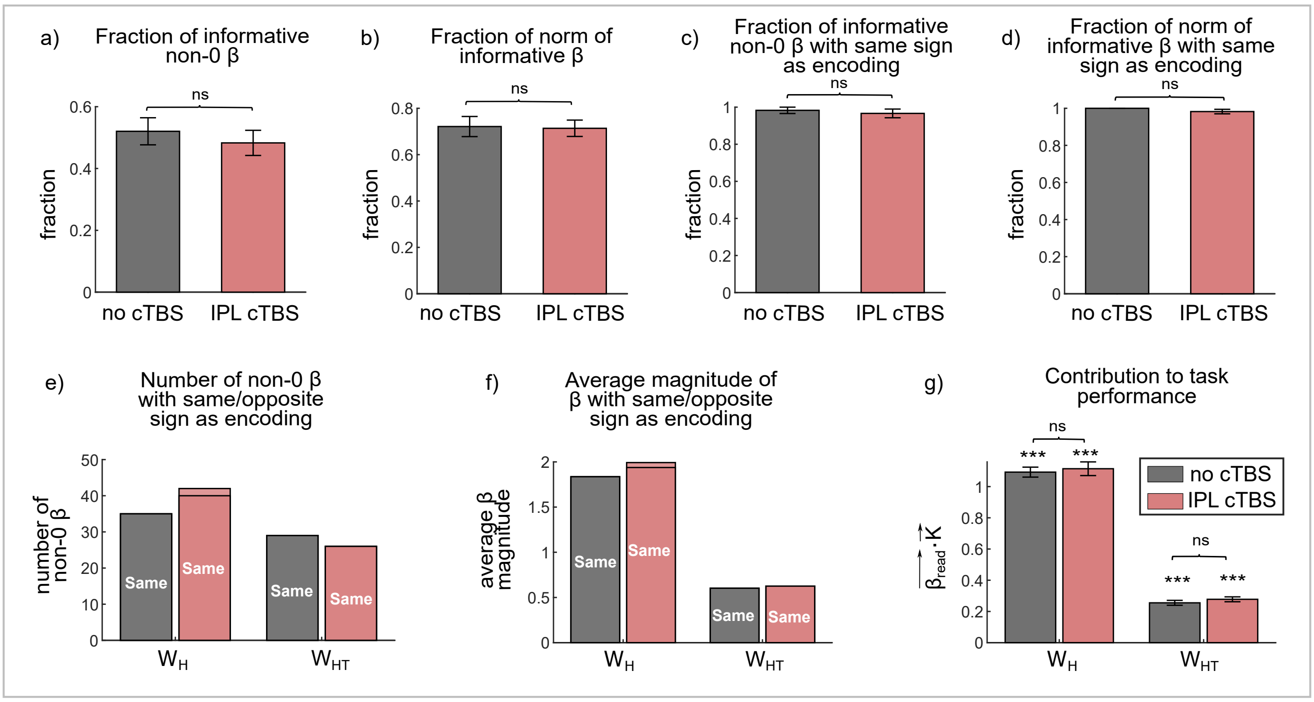


Supplementary Figure 6

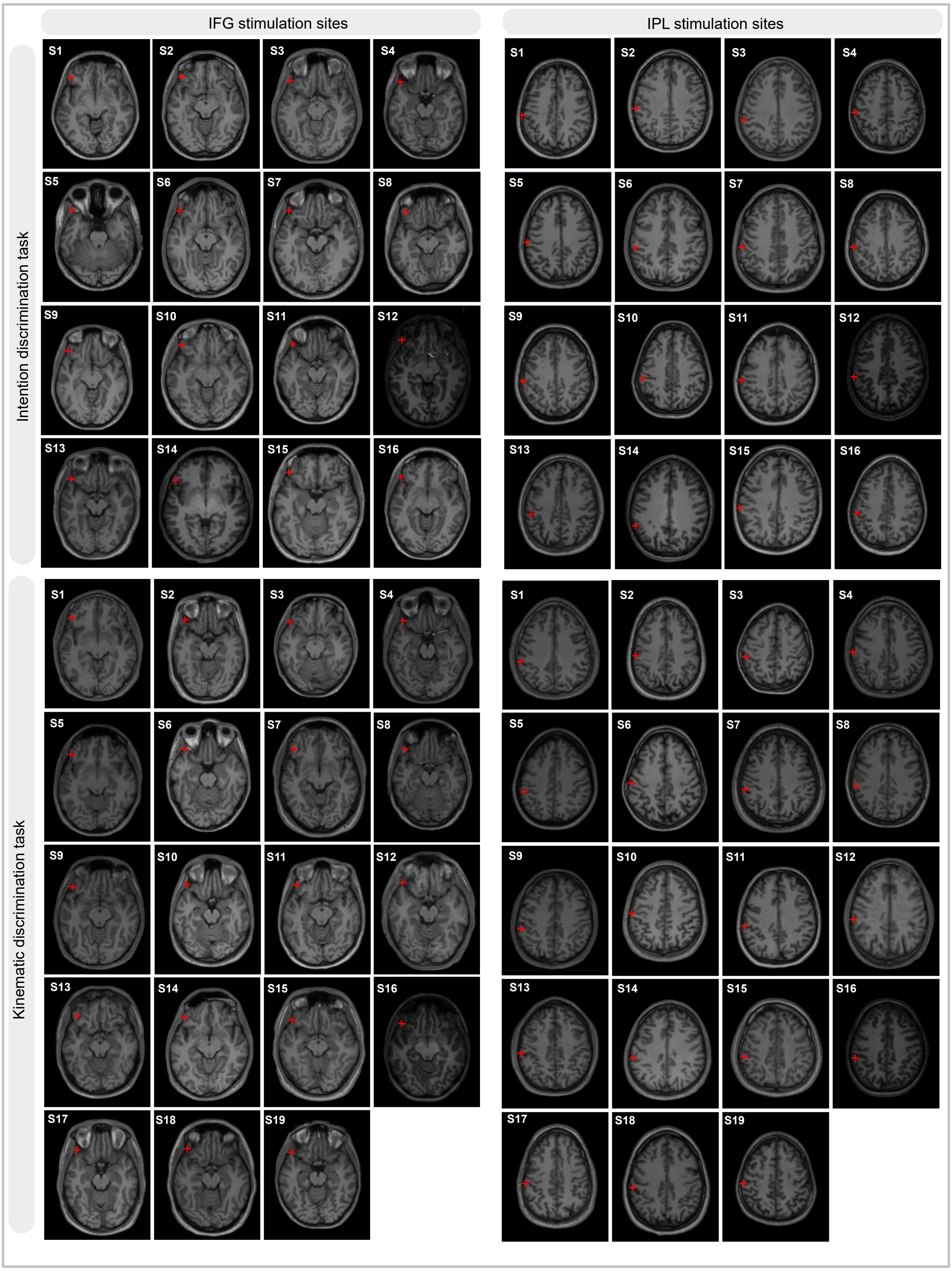


Supplementary Figure 7
